## Supplementary figures and images for "Transcription factors in the fungus *Aspergillus nidulans*: Markers of genetic innovation, network rewiring and conflict between genomics and transcriptomics"

### Figure S1

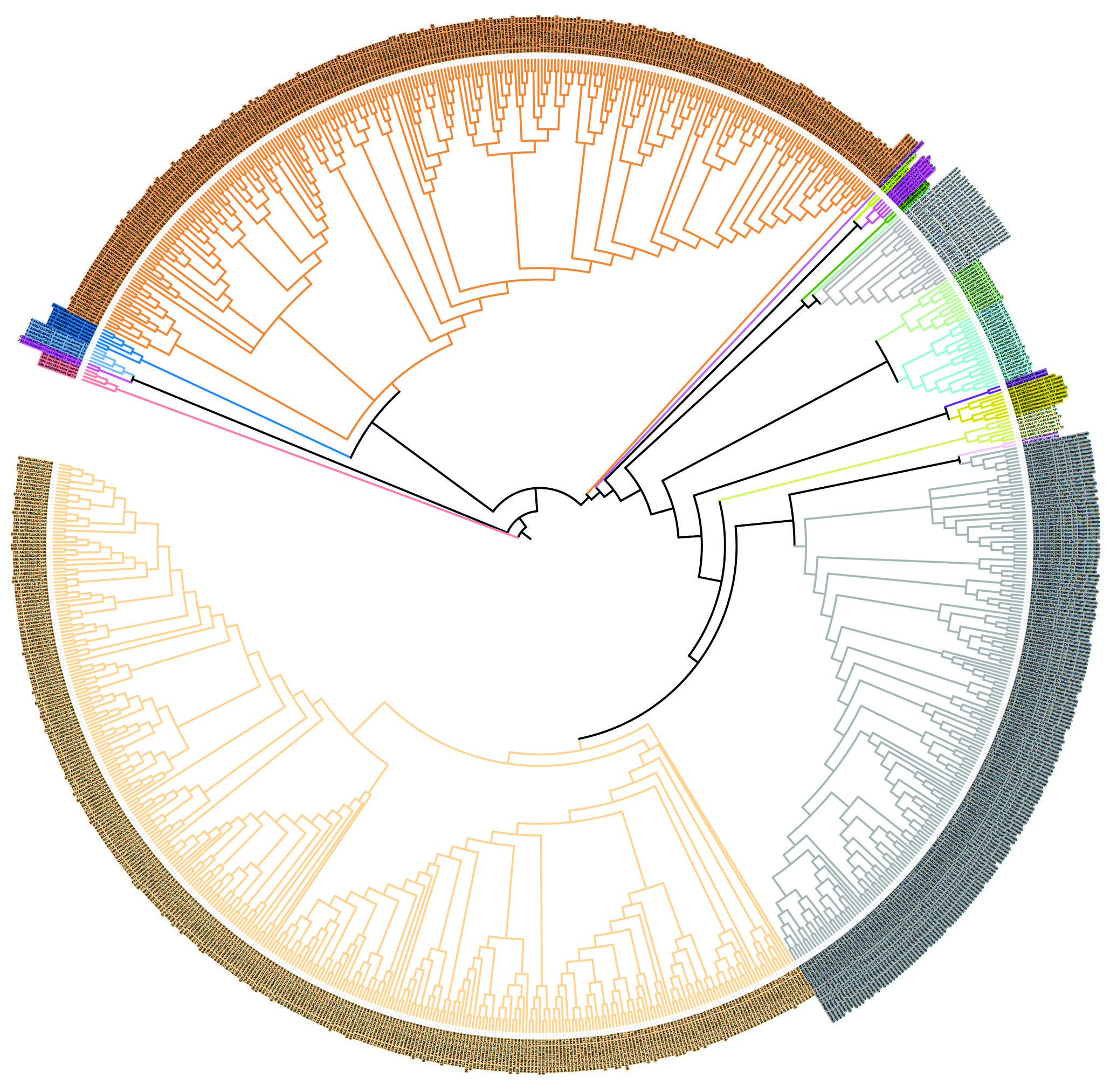

### Figure S2

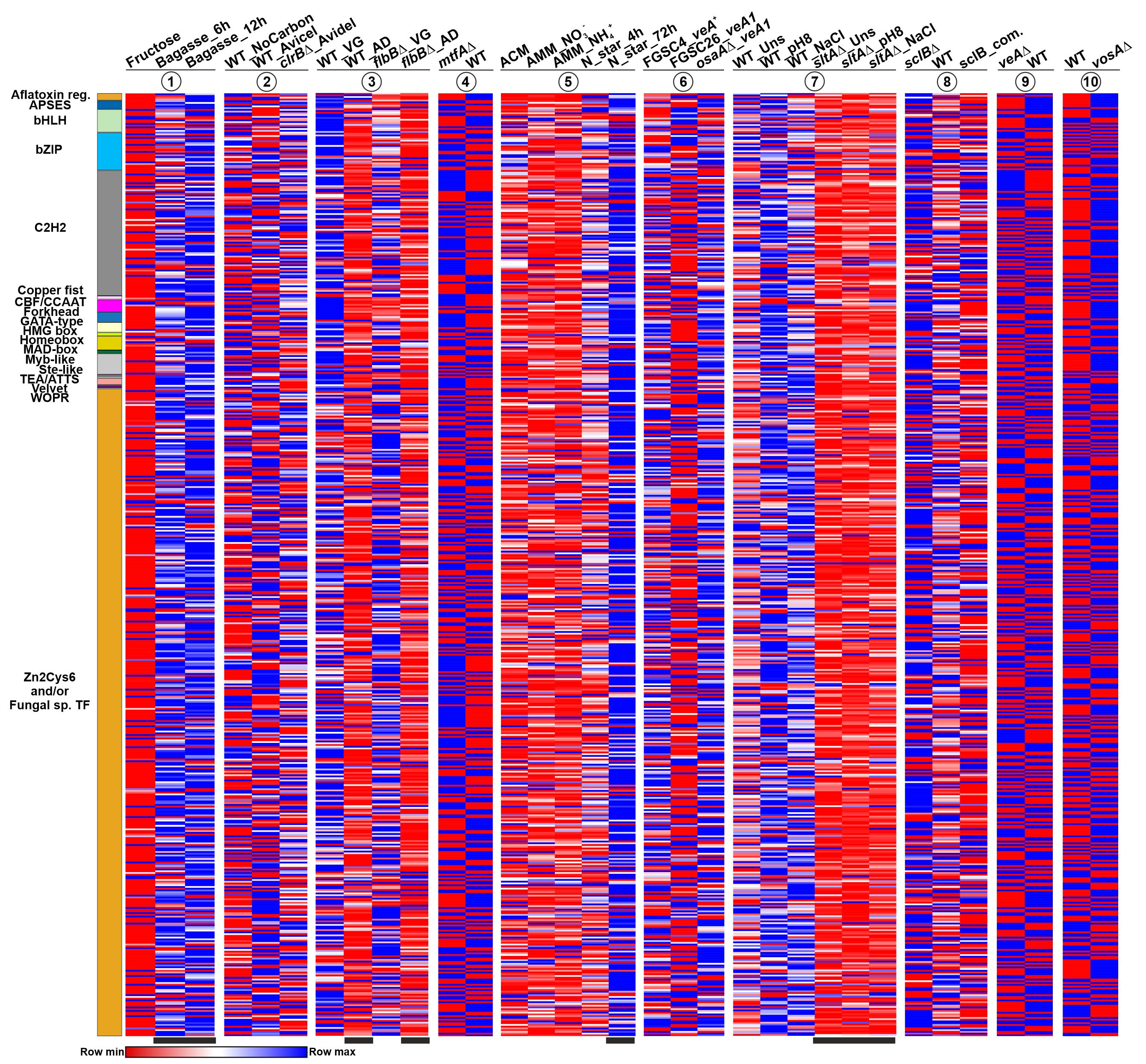

### Figure S3

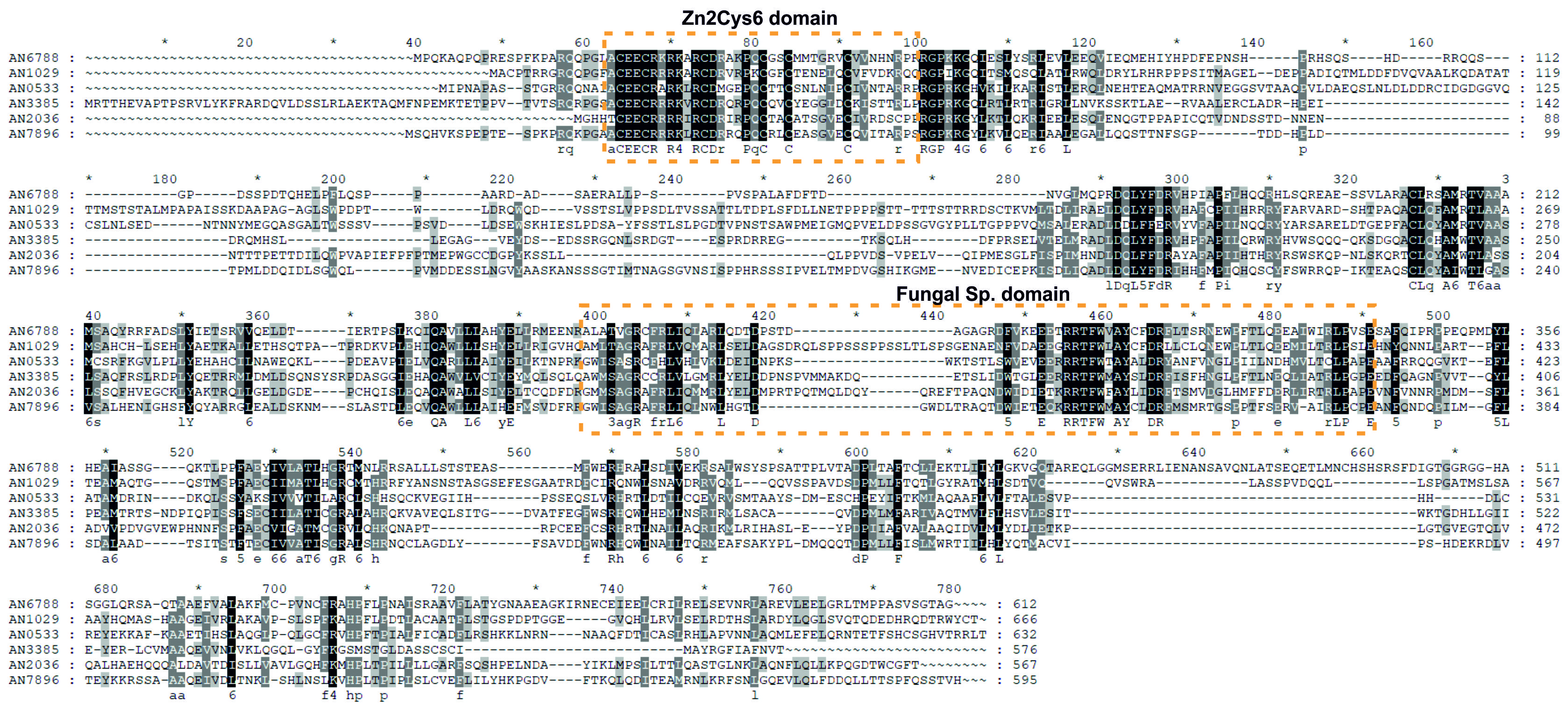

### Figure S4

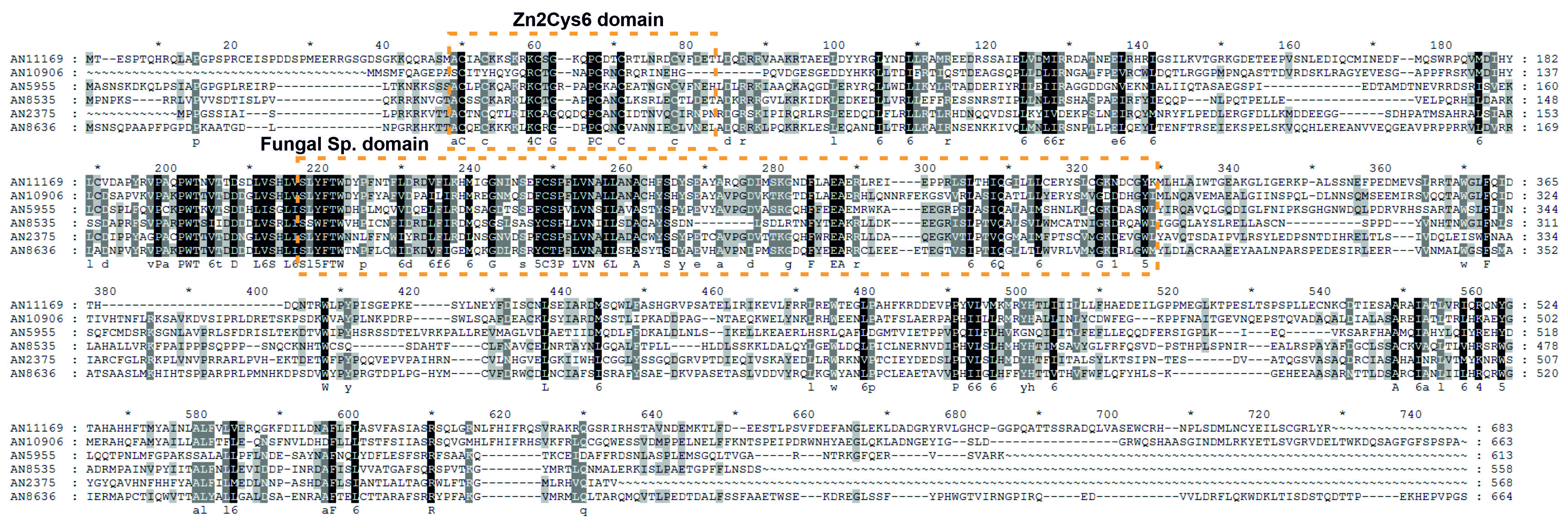

### Figure S5

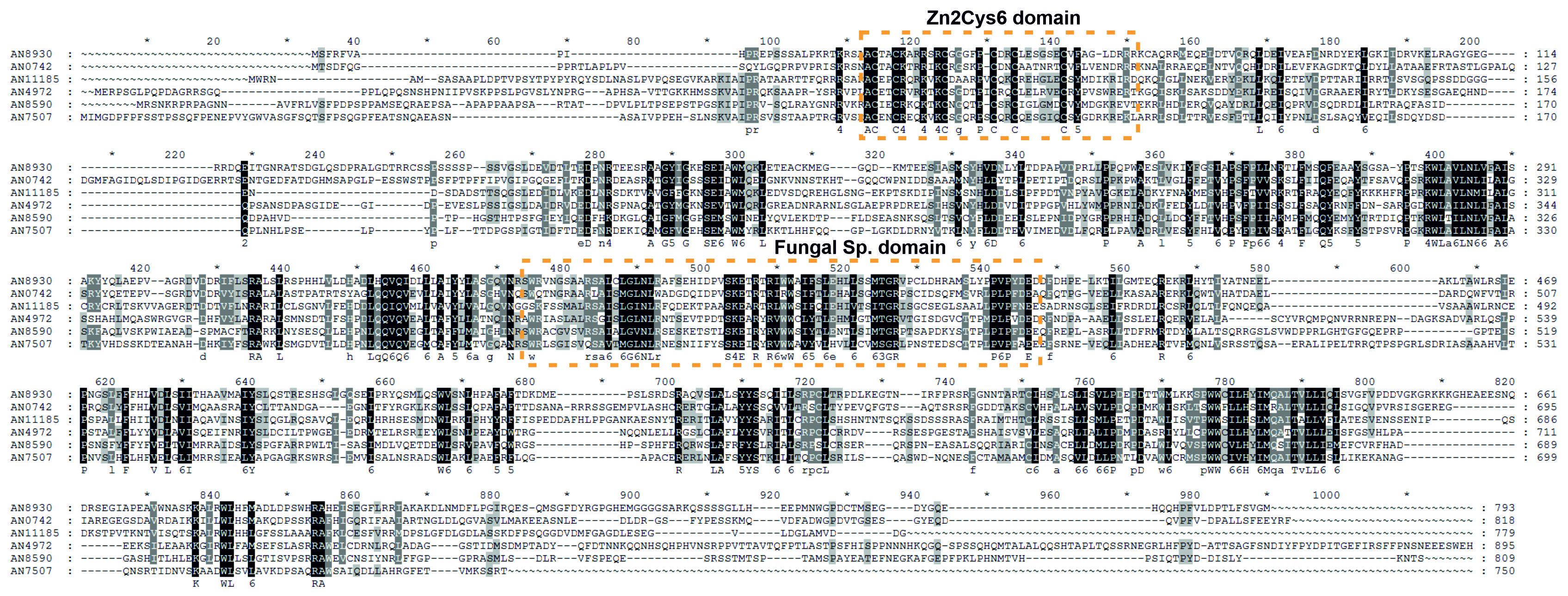

### Figure S6

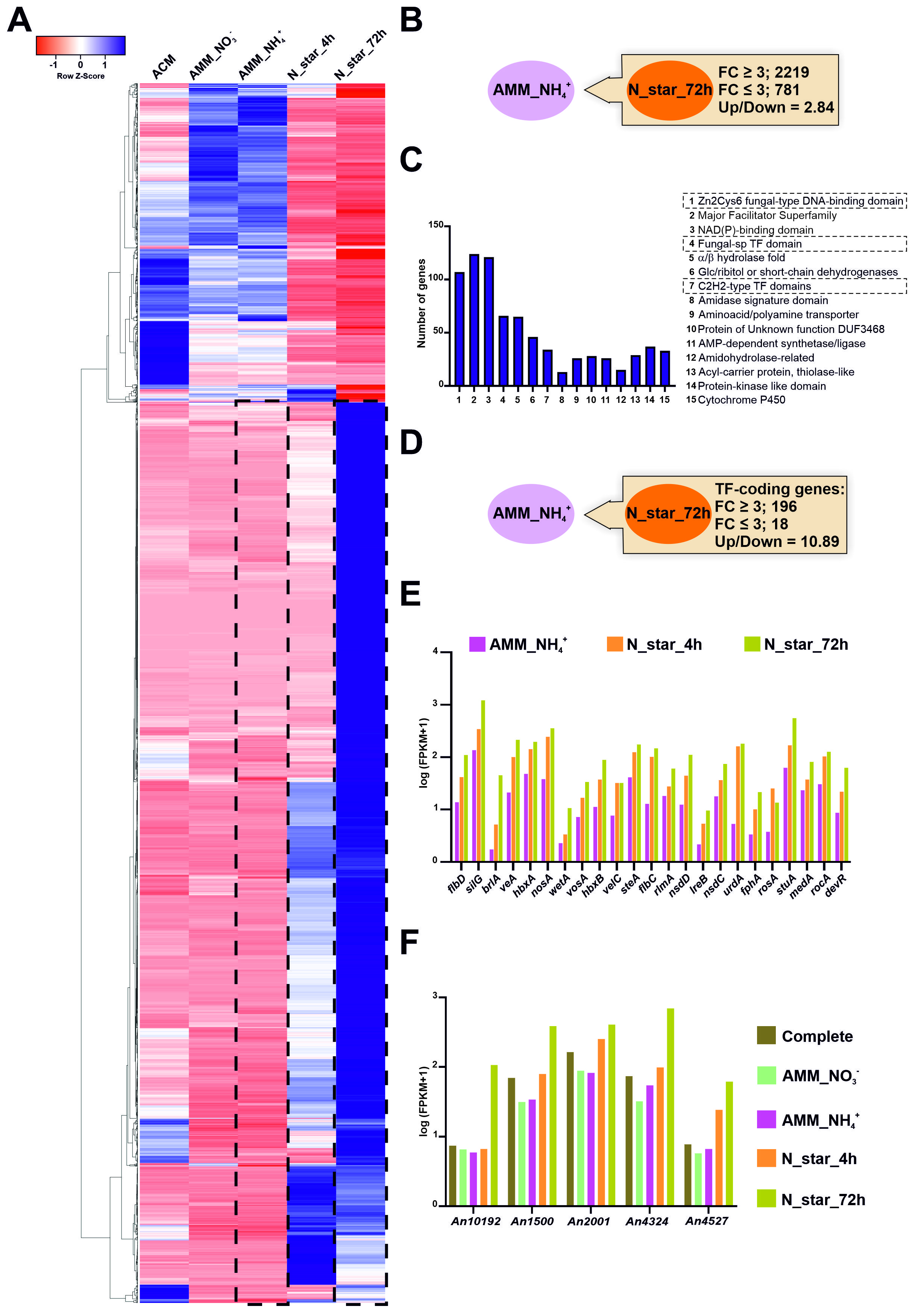

### Figure S7

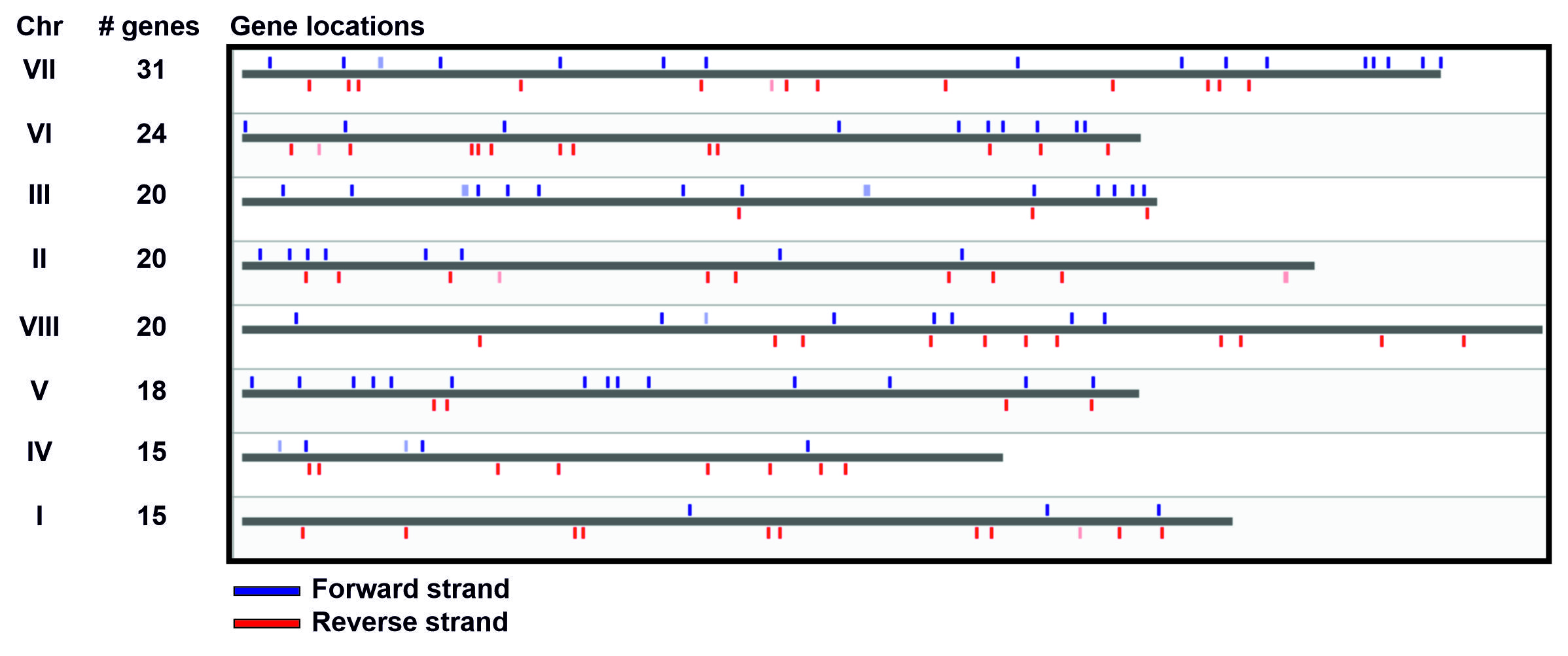
