## Supplementary material for "Transcription factors in the fungus *Aspergillus nidulans*: Markers of genetic innovation, network rewiring and conflict between genomics and transcriptomics": Table S1

**Table S1: Oligonucleotides used in this study.**

| **Name** | **Sequence (5´-3´)** |
| --- | --- |
| BrlA-PP1 | CAGCCGGGTACTGGAAGCACC |
| BrlA-GSP2 | TTCATCCCAGCCGTCCAGGC |
| BrlA-GFP1 | GCCTGGACGGCTGGGATGAAGGAGCTGGTGCAGGCGCTGGAGCC |
| BrlAGFP2 | AGATCAGCCCTCTTTGTTTCTGTTTCAGTCTGAGAGGAGGCACTGATGCG |
| BrlA-GSP3 | TGAAACAGAAACAAAGAGGGCTGATCT |
| BrlA-GSP4 | CCGCTTCCTACCCCGAATGG |
| BrlA-ZnFDw | CTGCACCTGCTTGATGACCTGTGG |
| MsnAZnFforBrlAUp | CGATTGCGACCACCCCGTTTGCTGTGAGTCCTGGCGTGCTGGGCAAGG |
| MsnAZnFforBrlADw | CCACAGGTCATCAAGCAGGTGCAGACATTCGTCTGCAATCTTTGCTCCCGC |
| BrlA-ZnFUp | AGCAAACGGGGTGGTCGCAATCG |
| BrlAp2Dw | GAGGAAGTGGTAAACTGGCGGATGG |
| BrlA±(+1) | GAAGATCTCGCCGCTCCTCCTCC |
| An10192-PP1 | CGACCAAAGAAGCCGCGATCACAGG |
| An10192-PP2 | CATTGGCGTTGTCTGATTCAAGTGTGTCTTCG |
| An10192-GSP3 | TAACTCTTATTGATGCGCCTAGCTGTGTTCTTTGAGC |
| An10192-GSP4 | CGAAGGACTCTACGTAGACCTCGTGGGC |
| An10192-SMP1 | CGAAGACACACTTGAATCAGACAACGCCAATGACCGGTCGCCTCAAACAATGCTCT |
| An10192-GFP2 | GCTCAAAGAACACAGCTAGGCGCATCAATAAGAGTTAGTCTGAGAGGAGGCACTGATGCG |
| An10192-sPP1 | CCACTGCCCCTGGTTCCAAGAAAGC |
| An10192-sGSP4 | GGCCGCTAACACTACCCAATATCACACGC |
| An1500-PP1 | CGTTTCGTATTGGAGGTATGGCAAGCG |
| An1500-PP2 | CATCATTTTGACGGTGTTGGATCGC |
| An1500-GSP3 | GATGGCTTGATTTGCGAGCTGGAATTGG |
| An1500-GSP4 | GCATCTGCAATGACATGTCCCTCACC |
| An1500-SMP1 | GCGATCCAACACCGTCAAAATGATGACCGGTCGCCTCAAACAATGCTCT |
| An1500-GFP2 | CCAATTCCAGCTCGCAAATCAAGCCATCGTCTGAGAGGAGGCACTGATGCG |
| An1500-sPP1 | GTCAAAGCACAATTGGTGATGGCACG |
| An1500-sGSP4 | CAGCGTGATAAAGGAGGCAATTCGAAGC |
| An2001-PP1 | GAAGCTAGTAGTGCAGCGTCAGCCACC |
| An2001-PP2 | CATGGTCACTGTAGCCTATATCCAGAGATAAAGACAAC |
| An2001-GSP3 | GCGAACTAGAACGACTGAATCTAAGCACATATATATGGC |
| An2001-GSP4 | GGATCTTTTGCGGCAGGCACC |
| An2001-SMP1 | GTTGTCTTTATCTCTGGATATAGGCTACAGTGACCATGACCGGTCGCCTCAAACAATGCTCT |
| An2001-GFP2 | GCCATATATATGTGCTTAGATTCAGTCGTTCTAGTTCGCGTCTGAGAGGAGGCACTGATGCG |
| An2001-sPP1 | CTGCGGAAAGGGCACTTACTTGCG |
| An2001-sGSP4 | GGAGGCGAAGCAACTTAGTCAGTCGG |
| An4324-PP1 | CTCGCCGCAATTGAGAGTCACCG |
| An4324-PP2 | CATTGTGGACGGTGGTAAATATGTACGAGAAG |
| An4324-GSP3 | TAACGCATTCTCTCTTTCTTCTCGCCAGATGC |
| An4324-GSP4 | CGAATCCGAATCCGTCTCCGAACCG |
| An4324-SMP1 | CTTCTCGTACATATTTACCACCGTCCACAATGACCGGTCGCCTCAAACAATGCTCT |
| An4324-GFP2 | GCATCTGGCGAGAAGAAAGAGAGAATGCGTTAGTCTGAGAGGAGGCACTGATGCG |
| An4324-sPP1 | GATGCGCGAGGTCATTCGGGACG |
| An4324-sGSP4 | CATCCGTATAAGGCAAGCTGCGTCACG |
| An4527-PP1 | CATTGACTGCTTGAAGGGAGATGTGACC |
| An4527-PP2 | CATATTGGTCAATGTGCAAGCTTTCCAATCACG |
| An4527-GSP3 | TAGATATCGAACGGCTGTGACGACGTAACG |
| An4527-GSP4 | CCCACAGCCACAGATAGACTTCCATTCG |
| An4527-SMP1 | CGTGATTGGAAAGCTTGCACATTGACCAATATGACCGGTCGCCTCAAACAATGCTCT |
| An4527-GFP2 | CGTTACGTCGTCACAGCCGTTCGATATCTAGTCTGAGAGGAGGCACTGATGCG |
| An4527-sPP1 | GTGAGCTTACCAAGTGCCCGACCTGC |
| An4527-sGSP4 | CTGATGACGATAATCTGGGGTAGGAAGGC |
